# Environmentally-induced DNA methylation is inherited across generations in an aquatic keystone species (*Daphnia magna*)

**DOI:** 10.1101/2021.12.05.471257

**Authors:** Nathalie Feiner, Reinder Radersma, Louella Vasquez, Markus Ringnér, Björn Nystedt, Amanda Raine, Elmar W. Tobi, Bastiaan T. Heijmans, Tobias Uller

## Abstract

Environmental stress can result in epigenetic modifications that are passed down several generations. Such epigenetic inheritance can have significant impact on eco-evolutionary dynamics, but the phenomenon remains controversial in ecological model systems. Here, we used whole-genome bisulfite sequencing on individual water fleas (*Daphnia magna*) to assess whether environmentally-induced DNA methylation can persist for up to four generations. Genetically identical females were exposed to a control treatment, one of three natural stressors (high temperature, zinc, microcystin), or the methylation-inhibitor 5-azacytidine. After exposure, lines were propagated clonally for four generations under control conditions. We identified between 70 and 225 differentially methylated CpG positions (DMPs) between controls and F1 individuals whose mothers (and therefore they themselves as germ cells) were exposed to one of the three natural stressors. Between 46% and 58% of these environmentally-induced DMPs persisted until generation F4 without attenuation in their magnitude of differential methylation. DMPs were enriched in exons and largely stressor-specific, suggesting a possible role in environment-dependent gene regulation. In contrast, treatment with the compound 5-azacytidine demonstrated that pervasive hypo-methylation upon exposure is reset almost completely after a single generation. These results suggest that environmentally-induced DNA methylation is non-random and stably inherited across generations in *Daphnia*, making epigenetic inheritance a putative factor in the eco-evolutionary dynamics of fresh-water communities.

**Author summary:** Water fleas are important keystone species mediating eco-evolutionary dynamics in lakes and ponds. It is currently an open question in how far epigenetic inheritance contributes to the ability of *Daphnia* populations to adapt to environmental stress. Using a range of naturally occurring stressors and a multi-generational design, we show that environmentally-induced DNA methylation variants are stably inherited for at least four generations in *Daphnia magna*. The induced variation in DNA methylation are stressor-specific and almost exclusively found in exons, bearing the signatures of functional adaptations. Our findings imply that ecological adaptations of *Daphnia* to seasonal fluctuations can be underpinned by epigenetic inheritance of DNA methylation without changes in gene frequencies.

## Introduction

Environmental stress can cause systemic changes in development and physiology. Such changes have been shown to occasionally span several generations [e.g., 1, 2, 3]. Many of the responses involve changes in the molecular machinery that is associated with DNA and contributes to gene regulation. DNA methylation is one of the most well studied epigenetic mechanisms in this context, but its involvement in transgenerational effects remains controversial in animals [4–7]. In mammals, inheritance of environmentally induced DNA methylation is limited by the fact that epigenetic marks are typically reset during reproduction [8–10], and this may explain why transgenerational persistence of environmentally induced DNA methylation appears rather uncommon [5]. In invertebrates, the transgenerational persistence of stochastic or environmentally induced DNA methylation variation is poorly studied. This is partly because DNA methylation is of limited significance in traditional model systems [e.g., Drosophila; 11]. However, recent studies suggest that environmentally induced variation in DNA methylation can be passed on to subsequent generations in insects, and perhaps other invertebrates as well [12–14].

Water fleas of the genus *Daphnia* are common in lakes and ponds, where they play central roles in the functioning of ecological interactions, food webs, and nutrient cycling [15]. How *Daphnia* respond to environmental change can have strong impact on community and ecosystem dynamics, making *Daphnia* a model system to understand the interactions between phenotypic plasticity, adaptive evolution, and ecology on contemporary time scales [16]. Such eco-evolutionary dynamics may be fundamentally altered if environmentally induced responses are inherited, for example, via epigenetic mechanisms [17]. However, the extent and specificity of transgenerational persistence of environmentally induced epigenetic variation remains poorly understood, not only in *Daphnia* but in ecological model systems in general [18, 19].

In addition to its role as a keystone species, there are several others reasons why *Daphnia* is particularly useful to study epigenetic inheritance. Individuals frequently reproduce clonally, which makes it possible to study epigenetic inheritance without the confounding effects of genetic variation [20, 21]. Furthermore, *Daphnia* inhabit waters with seasonal environmental variation, spanning periods of multiple asexual generations, a situation that should favour incomplete epigenetic resetting [22–24]. Thus, *Daphnia* may be particularly likely to have evolved mechanisms that enable context- and gene-specific inheritance of gene regulation. Since *Daphnia* are carrying their offspring in an actively ventilated brood pouch, maternal exposure to environmental stressors can have effects on future generations by directly affecting embryos or germ cells, similar to the situation in mammals. Such effects on offspring phenotype and fitness are commonly observed and occasionally carry over to more than two generations (e.g., UV [25], microcystin [26], temperature [27], and predator cues [28]). A candidate epigenetic mechanism underlying transgenerational plasticity in *Daphnia* is DNA methylation. Despite that DNA methylation in water fleas is occurring at low levels [∼0.6%; 29, 30], it is typically enriched in exons and is positively correlated with levels of gene expression [29]. There is also some evidence that environmentally induced variation in DNA methylation can be passed on to subsequent generations [31–33]. However, a rigorous assessment of genome-wide patterns of inheritance on the individual level has not been performed to date.

In this study, we chose four stressors that have been shown to affect global DNA methylation. Three are stressors that *Daphnia* encounter in the wild (*Microcystis aeruginosa*, a cyanobacteria producing the toxin microcystin, zinc, and elevated temperature) and one is a toxin not naturally encountered by *Daphnia* (5-azacytidine, a compound that inhibits the function of the DNA methyltransferase DNMT1 and thereby causes hypo-methylation [34]). Using a multi-generational experimental design (Figure 1), we explored (1) the immediate impact of the stressors on genome-wide DNA methylation levels in F1 individuals, which were germ cells during maternal exposure, by identifying differentially methylated cytosine positions (DMPs), (2) whether these DMPs are specific for each stressor, (3) whether DMPs persist across four generations and (4) the putative biological function of these DMPs.

**Figure 1.**
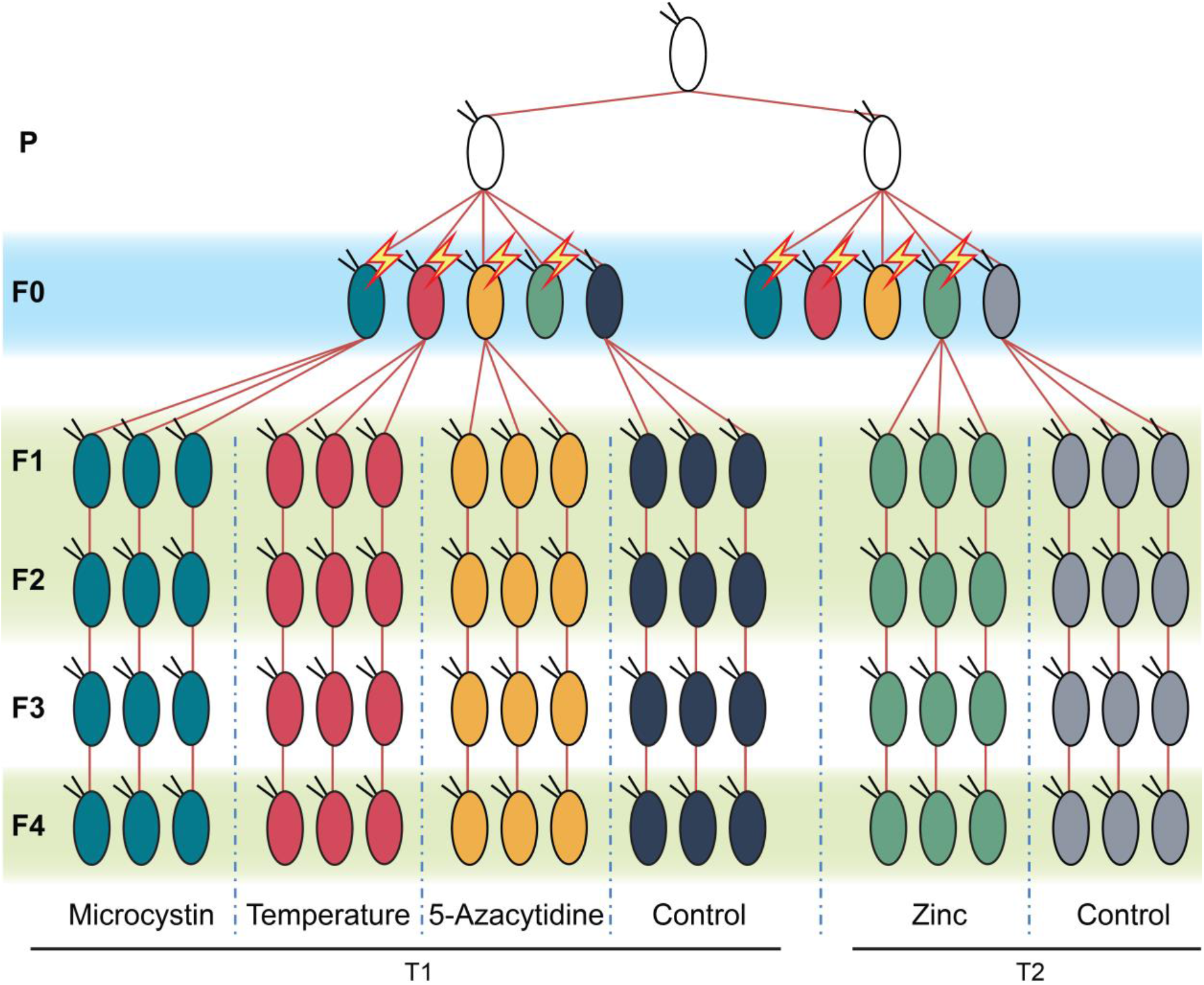
Schematic representation of the experimental design. Clonal siblings (generation P) were divided into two lines (experiment T1 and T2) and their offspring (F0) were exposed to environmental stressors (microcystin, high temperature, 5-azacytidine, or zinc) or kept under control conditions (one per line). This experiment was replicated twice (T1 and T2) and run in parallel to account for potential incompleteness due to the extinction of maternal lines. The most complete experiment for each stressor was subjected to further analyses (as indicated, mortality in the zinc-exposed line resulted in incomplete data and the analyses for zinc therefore make use of data from the second experiment). Individual *Daphnia* of the generation F0 were exposed to environmental stressors from birth to first reproduction (more detail in Table S1). Since the maternal treatment stopped before the egg cells were released into the brood pouch, the F1 generation was exposed as germ cells to the stressors but not as embryos. Five offspring per exposed (or control) mother were selected and allowed to propagate until generation F4 under control conditions. Red lines represent propagation of second brood offspring. Individual *Daphnia* of each treatment group and generation F1, F2 and F4 were subjected to whole-genome bisulfite sequencing (indicated by light green boxes). Each experimental unit included in the final analysis consisted of three individuals, except for T1-5-azacytidine-F1 and T1-Control-F4, which consisted of two replicate individuals, and T2-control-F1 and T1-control-F2, which consisted of four replicate individuals. Tests of transgenerational persistence of environmentally induced DNA methylation were analysed separately for T1 and T2. For more details, see Methods and Tables S1 and S2.

## Results

### Environmental stressors negatively affect the reproductive output of exposed *Daphnia*

Fitness assays on the reproductive output of individuals showed a direct effect on the stressor-exposed F0 generation relative to control samples for all stressors except elevated temperature (Figure 2). These negative fitness effects were most pronounced for *Daphnia* exposed to toxic cyanobacteria (microcystin treatment). However, this treatment line, as well as the zinc treatment line, regained reproductive fitness already in the F1 generation. The negative fitness effect of the 5-azacytidine treatment persisted until generation F3. Elevated temperature negatively affected reproductive fitness in generations F1 and F3 (Figure 2). Thus, the treatments significantly affected reproductive output of exposed individuals and caused maternal and transgenerational effects on fitness that demonstrate the physiological relevance of the stressors.

**Figure 2.**
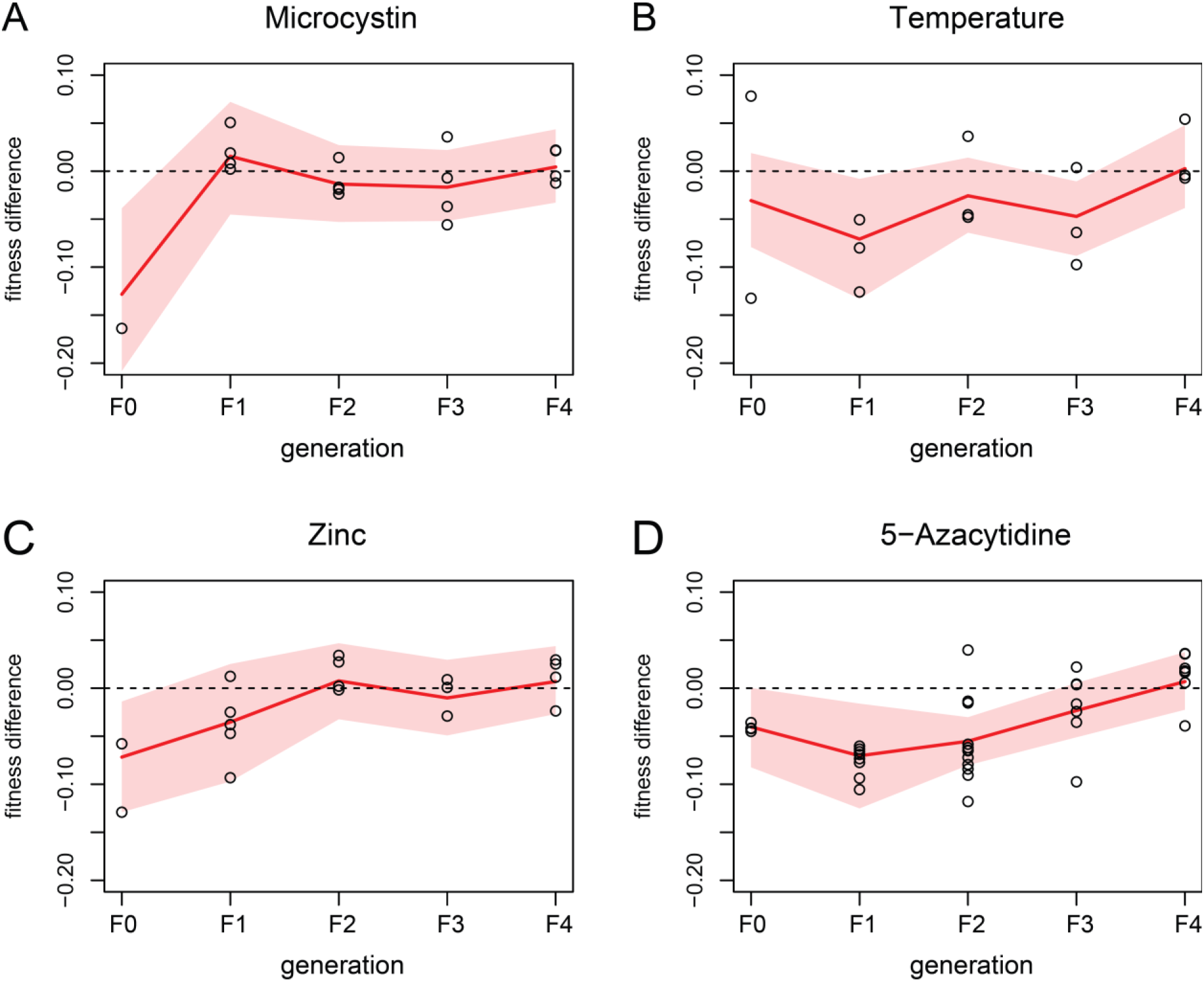
Fitness effects of the environmental stressors. (A-D) Each plot shows the fitness difference of exposed *Daphnia* per generation relative to the control conditions (marked as dashed line). Fitness estimates refer to lifetime reproductive output derived from the age at first and second reproduction and the size of the first and second brood (see Methods). Red lines mark the means and shaded areas the 95% credible intervals. Observed values (statistically corrected for clone line effects) are plotted as black circles. Note that fitness data was recorded for all individuals included in this study, including those not selected for bisulfite sequencing. For (D) 5-azacytidine the fitness effects were sustained until generation F2, whereas for (A) microcystin and (C) zinc the effects disappeared after generation F1. For (B) high temperature, fitness effects lasted for several generations (until generation F3), though fitness was significantly different from the control samples only for generation F1 and F3.

### Genome-wide methylation levels are consistently low and reduced by 5-azacytidine

Consistent with previous studies in *D. magna* [0.74%; 29] [0.52%; 30], we found an overall low proportion of CpG sites in a methylated state (Figure 3). In control samples, 0.50% (SD: 0.02%) of all CpG sites were methylated, and similar proportions were found in *Daphnia* that were exposed to one of the natural stressors (high temperature, zinc or microcystin) as germ cells (F1), and in their non-exposed descendants (F2 and F4). As expected, the stressor 5-azacytidine, which inhibits DNA methylation, caused a tenfold decrease in methylation levels of CpG sites in the generation F1 (0.05%; SD: 0.02%, *P*-value <0.01) relative to control F1. Subsequent generations (F2 and F4) of 5-azacytidine exposed *Daphnia* showed CpG methylation levels that approached the levels of the control samples, but remained at a consistently lower level (Figure 3).

**Figure 3.**
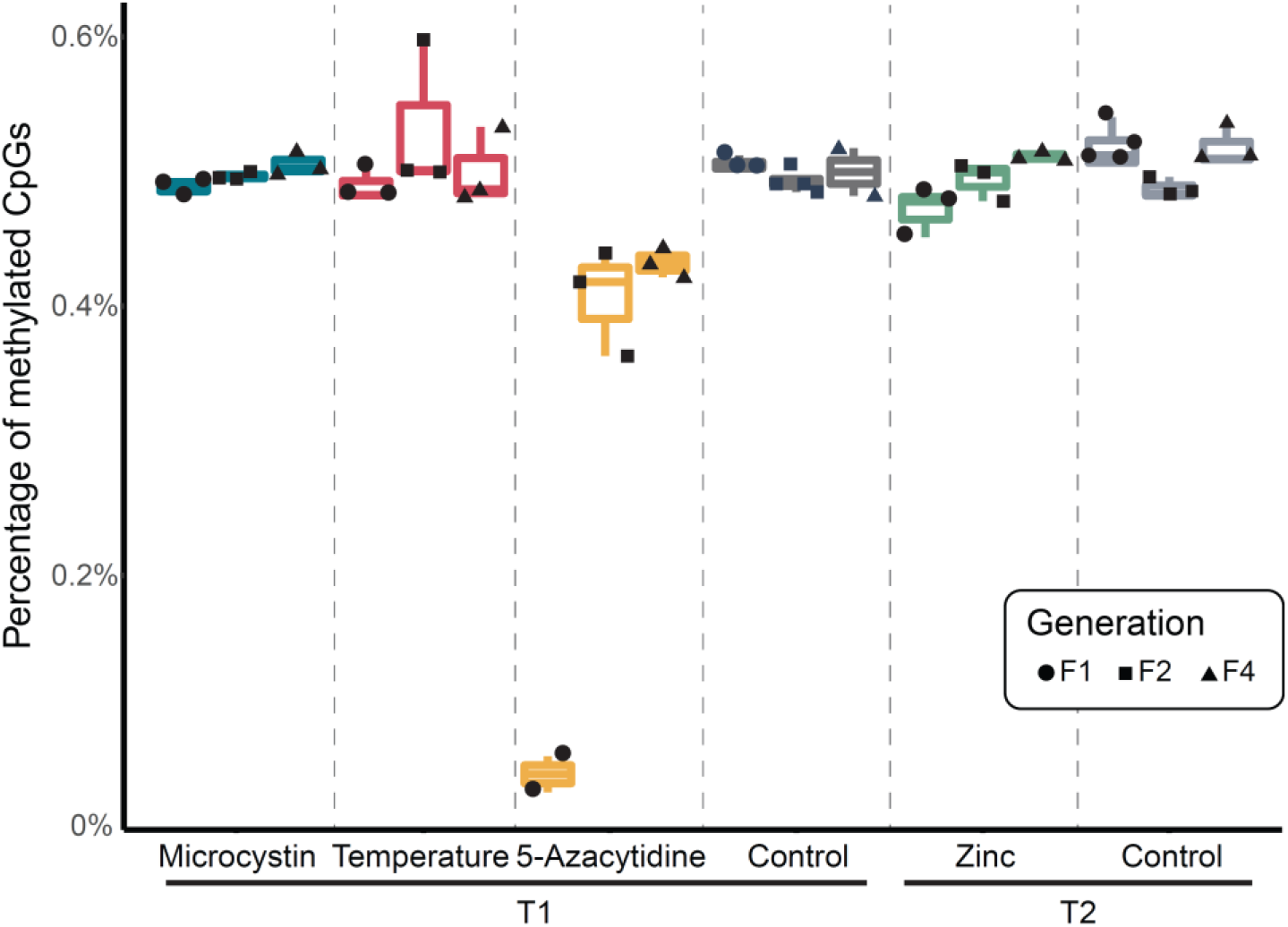
Overall levels of CpG methylation across samples. Plots show the percentages of CpGs that are methylated relative to the total number of CpGs in the genome for each treatment group. Boxes are coloured according to treatment group. The lower, median and upper hinges correspond to the first, second and third quartiles respectively. Whiskers indicate the range that lies within 1.5 times of the interquartile ranges. Black symbols indicate the individual data points according to generation.

### Germ-cell exposure to environmental stressors leads to differential methylation of CpG sites

We detected an effect on genome-wide patterns of methylation in individual F1 *Daphnia* that were exposed to natural stressors as germ cells. We identified 70 DMPs in the F1 generation exposed to thermal stress (relative to the nine control samples in the F1, F2 and F4 generations), 76 DMPs in *Daphnia* exposed to zinc, and 225 DMPs in *Daphnia* exposed to microcystin (at 5% FDR; Tables S3-S5).

Consistent with the strong signal of demethylation through 5-azacytidine, we found 2,231 DMPs in F1 *Daphnia* of this treatment line. While pairs of treatments shared a low number of DMPs (Table S6), we found no DMPs in F1 that were shared by all four stressor groups, and also no DMPs shared between the three natural stressors (thermal stress, zinc and microcystin). Thus, the induced methylation changes were largely stressor-specific.

### A large proportion of environmentally-induced DMPs persist until the F4 generation

After calling DMPs for each stressor and for each generation (at 5% FDR), we intersected DMPs across generations per stressor to assess their overlap and thus the persistence of the stress response. For example, of the 225 DMPs detected in F1 *Daphnia* from the microcystin treatment, 57.8% (130 sites) were also differentially methylated compared to control samples in generation F2 and F4 (Figure 4A; Table S7). For zinc and temperature, 53.9% and 45.7% of the environmentally induced methylation variants in F1 persisted until the F4 generation (Tables S8 and S9). Across the natural stressors (high temperature, zinc or microcystin), the number of DMPs shared among three generations (F1, F2 and F4) was greater than expected by chance (*P*-value derived from 1000 permutations: <0.001 for all three stressors), and also greater than the number of DMPs unique to F2 or F4 (Figure 4A). These results demonstrate transgenerational stability of stress-induced methylation marks.

**Figure 4.**
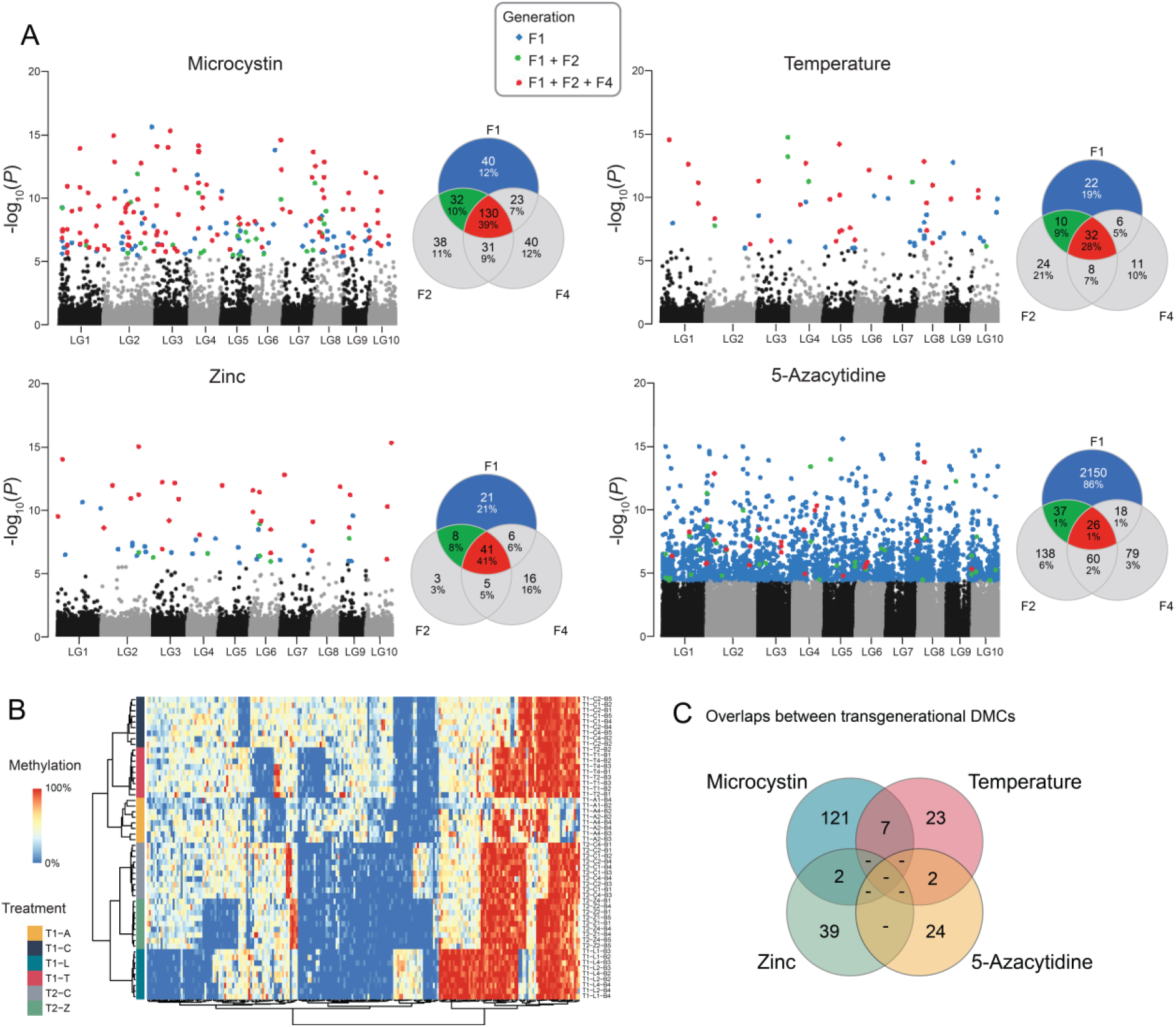
Transgenerational inheritance of DMPs. (A) Manhattan plots showing the genomic positions of DMPs (left) and Venn-diagrams showing the overlap of DMPs across generation F1, F2 and F4 (right) for each of the four stressors. DMPs are color-coded according to their transgenerational persistence as indicated. Numbers in the centre of the Venn-diagrams give the number of transgenerational DMPs. Percentages of all DMPs are provided below absolute numbers of DMPs in each group. (B) Heatmap of the methylation level for all 229 transgenerational DMPs across all samples included in this study. Label names consist of experiment (T1 or T2), treatment and generation (A, 5-azacytidine; L, microcystin; T, temperature; Z, zinc; followed by a number indicating the generation), and clonal line number (B1-B5). (C) Venn diagram presenting the overlap of transgenerational DMPs across the treatments. No DMP was shared across three or all four treatments, but between two and seven are shared by two stressors.

Since 5-azacytidine induced a strong hypomethylation in the F1 generation exposed as germ cells, followed by a re-methylation in following generations (Figure 3), the patterns of transgenerational inheritance were different from that of the other stressors (Figure 4B). Only 1% (26 sites) of the environmentally induced methylation variants in F1 persisted until the F4 generation, all of which remained hypomethylated (Figure 4A; Table S10). Thus, compared to the other three stressors, the number of DMPs within each generation were consistently higher, but their inter-generational overlap was lower.

To robustly verify that the treatment-induced transgenerational DMPs are stably persisting across generations and not due to stochastic events, we used two additional strategies for data analyses. Firstly, we applied permutations by randomly shuffling sample labels to generate a null hypothesis. These permutations demonstrated deflated *P*-values relative to the observed *P*-values (Figures S1 and S2) and produced no transgenerational DMPs except two in 100 permutations in the 5-azacytidine case. Secondly, to mitigate the potential bias stemming from using the same set of control samples (i.e., all nine control samples) in each statistical test, we used an alternative strategy that identified candidate DMPs in the F1 and subsequently tested the significance of those candidates in F2 and F4 (for details and results, see Methods section). Both additional strategies broadly confirmed the existence of environmentally induced DMPs that persist for at least four generations.

Consistent with the limited overlap of DMPs of F1 *Daphnia* between the four stressors, we also found that transgenerational DMPs are largely stressor-specific. However, pairs of treatments (e.g., temperature and microcystin treatments) shared up to 7 DMPs, which was more than expected by chance (*P*-value derived from 1000 permutations: <0.001; Figure 4C).

### Transgenerational DMPs retain a consistent methylation pattern and are almost exclusively located in gene bodies

For all transgenerational DMPs of a given stressor, the sign of differential methylation (i.e., hypo- or hypermethylated) was consistent across the generations. Moreover, when comparing the effect sizes of differential methylation relative to control samples for a given stressors, we found that those observed in the F1 and in the F4 generation are remarkably similar in magnitude, and no sign of attenuation in the F4 generation was observed (Figure 5A).

**Figure 5.**
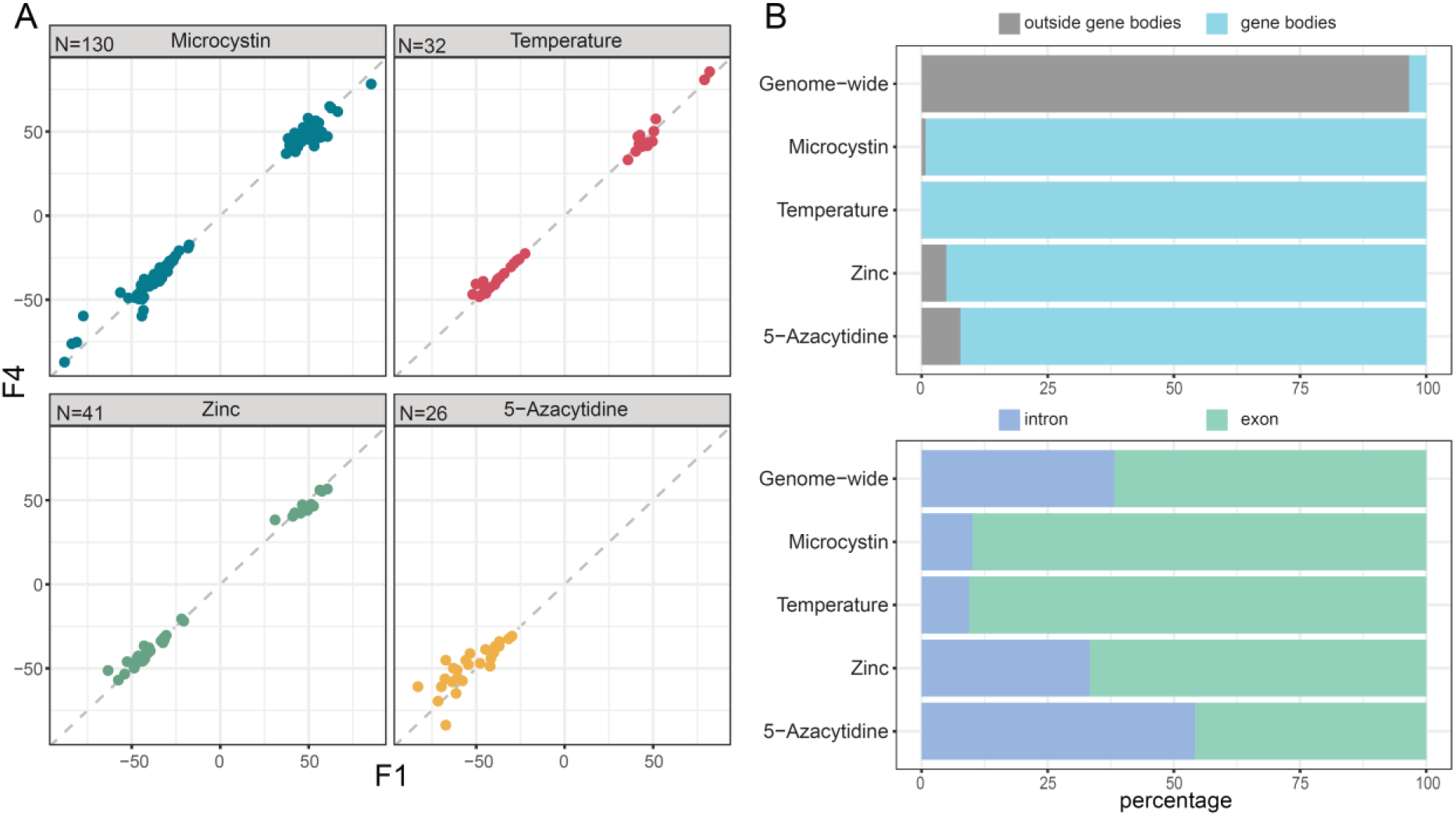
Characterization of transgenerational DMPs. (A) Effect sizes of the strength of differential methylation (methylation difference relative to the nine control samples) for the transgenerational DMPs of the F1 generation plotted against the corresponding effect sizes for the F4 generation. All DMPs lie close to the dashed line indicating equally strong effect sizes in the two generations, which shows that the effect sizes are consistent across generations. Note that all effect sizes for 5-azacytidine are negative, as expected, due to the hypomethylation caused by this compound. (B) Top panel shows the proportion of transgenerational DMPs that lie within gene bodies, and bottom panel further details the distribution of those to exons or introns. The comparison with the distribution of genome-wide CpG sites (N = 10,806,885) shows that the stressor-induced transgenerational DMPs are overrepresented in gene bodies, and tend to lie in exons rather than introns, except for the stressor 5-azacytidine.

To assign a putative functional role to the identified DMPs, we systematically characterized the genomic positions for two groups of DMPs: those that occurred in the F1 generation after developmental exposure (as germ cell) to a stressor (direct DMPs) and a subset of these, namely those that were stably inherited until generation F4 (transgenerational DMPs). We found that both direct and transgenerational DMPs are predominantly found in gene bodies (direct: 96.1%; transgenerational: 98.7%; Tables S3-S10, Figure 5B). Of all DMPs occurring in gene bodies, the majority lies in exons rather than introns (direct: 87.70%; transgenerational: 85.81%; Tables S3-S10, Figure 5B). Roughly half of the DMP-containing gene bodies contained at least two DMPs (direct: 54%; transgenerational: 42%) in close proximity to each other (median distance in bp, direct: 17; transgenerational, 1).

### Transgenerational DMPs are occurring in genes that exhibit stressor-specific functions

Although we did not find any overlap in the exact genes containing transgenerational DMPs induced by different stressors, members of the 60S (large) ribosomal protein family are consistently hypomethylated following exposure to a natural stressor (temperature, zinc or microcystin). A more systematic assessment of functional overlap using GO enrichment analysis showed that transgenerational DMPs were significantly enriched for a number of different functions. We found between 7 and 15 significant GO terms of biological processes and the top ten of these terms are shown in Figure 6. None of the GO terms were shared between two or more stressors (Tables S11 and S12). This shows that DMPs are largely occurring in different sets of genes for each stressor, and that these genes show no functional similarity (i.e., not the same GO terms).

**Figure 6.**
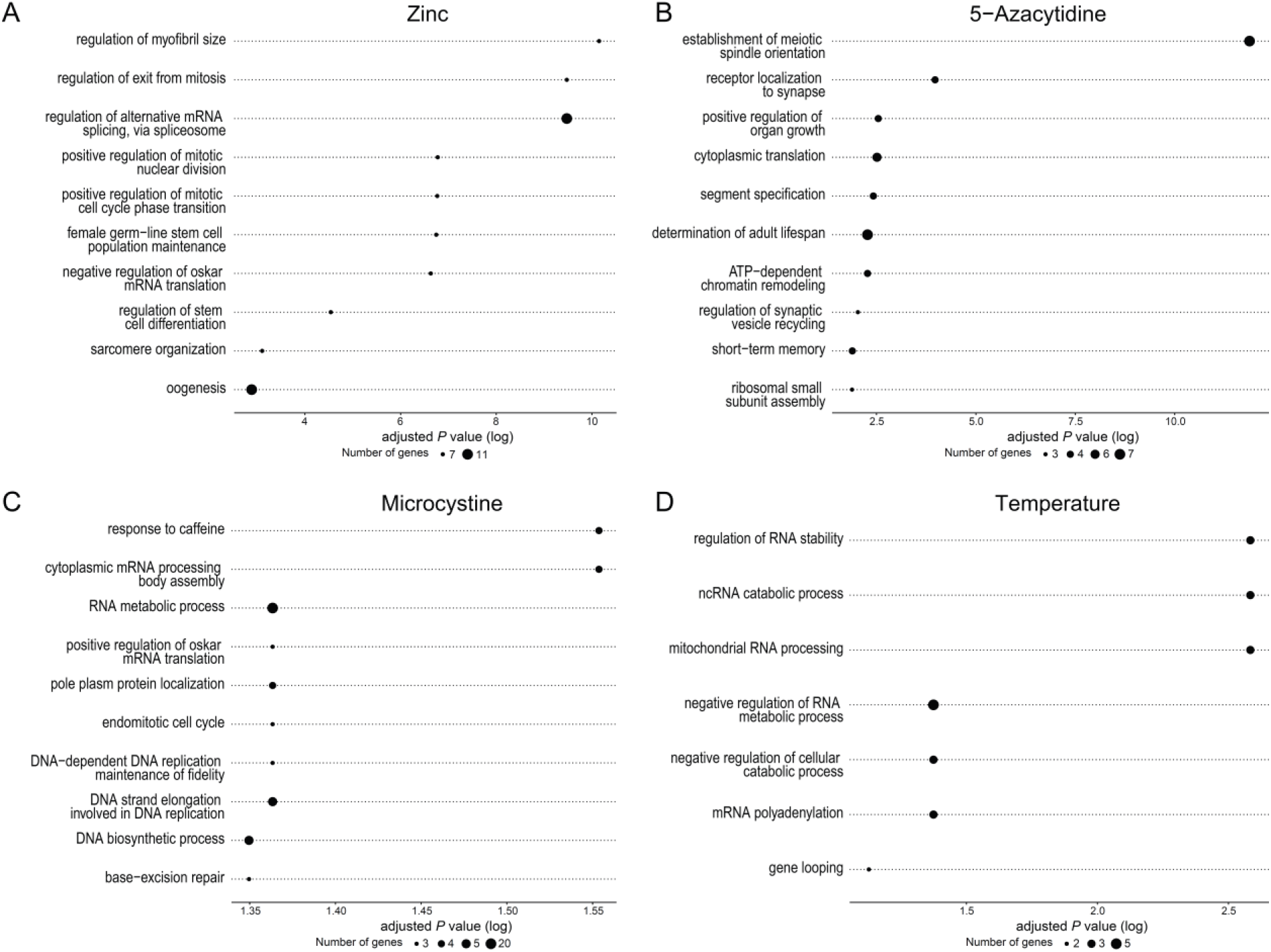
Putative functional role of transgenerational DMPs. (A-D) Top ten significantly enriched GO terms of biological processes for genes containing transgenerationally inherited DMPs across the four stressors. The size of each GO term represents the number of genes that it is represented by, and the position along the x-axis indicates its significance (log-transformed adjusted *P*-value).

### Candidate genes for stress responses reported in the literature are not differentially methylated

Finally, we cross-referenced genes identified as differentially methylated in our study with genes identified as differentially expressed upon stress exposure reported in the literature. We retrieved six studies that assessed gene expression differences in response to stressor exposure in *Daphnia sp.* We only included experimental designs that are comparable to the treatment conditions applied in our study, and that used genome-wide, unbiased approaches such as microarrays (four studies) or RNA-sequencing (two studies; Tables S13 and S14). Three of these studies used microcystin as stressor [35–37], two used zinc [38, 39], and one used temperature [40]. Between 6% and 15% of the key genes differentially expressed in response to microcystin, zinc, or temperature as stressor are highly functionally similar to genes identified as differentially methylated in our data, though gene identity was not identical (Tables S13 and S14). Most instances of genes identified as differentially expressed as well as differentially methylated concern genes with a well-described, broad functionality such as heat shock proteins or large ribosomal proteins (Tables S13 and S14).

The effects of 5-azacytidine on gene expression has not been assessed in a genome-wide, unbiased approach in *Daphnia*, but two studies reported the differential expression of several candidate genes (mostly DNA-methyltransferases, which are known to be the target of this drug) [41, 42]. We found that none of the key candidate genes differentially expressed in response to 5-azacytidine treatment was identified as differentially methylated in our analyses.

## Discussion

Despite extensive interest in epigenetic inheritance, the extent to which environmentally induced epigenetic marks are heritable in animals remains an open question. Here, we show that, in *Daphnia*, environmentally induced variation in DNA methylation in germ cells are specific to the stressor, and are stably inherited for at least four generations.

Maternal exposure to the natural stressors microcystin, zinc, and high temperature caused stressor-specific DNA methylation changes in their F1 offspring. Since the maternal treatment stopped before the egg cells were released into the brood pouch, the DMPs in F1 were likely induced in germ cells and persisted during cell differentiation to be evident in most, or all, cell types (and thus possible to detect by bisulfite sequencing of whole individuals; although differences in the relative number of cell types between treatments may also contribute). As expected, 5-azacytidine led to genome-wide hypomethylation, while the other stressors induced both hypo- and hypermethylation. One of the two genes that were affected in all three of the naturally occurring stressors – microcystin, zinc, and high temperature – was a 60S (large) ribosomal protein, which was consistently hypomethylated. Large ribosomal proteins have been repeatedly reported as being implemented in stress responses of a variety of organisms [43, 44], including salinity stress in *Daphnia* [32]. However, in general, the specific DMPs, the genes they reside in, and the putative functions of those genes (i.e., GO terms) were largely specific to each stressor.

The majority of environmentally induced DMPs were located in gene bodies, more precisely in exons, which is consistent with how DNA methylation appears to regulate gene expression in invertebrates [45, 46]. Indeed, some genes with DMPs do have putative functions for responding to these stressors, but few of the *a priori* candidate genes or pathways were identified as being differentially methylated. For example, DNA-methyltransferases, which are strongly down-regulated upon exposure to 5-azacytidine [41, 42], showed no signs of changes in methylation levels. Similarly, ABC transporter genes and nucleoside transporters, which are strongly expressed in response to microcystin [36] and zinc [38], respectively, were not differentially methylated. This might be explained by the fact that the differential expression of these candidate genes tends to be restricted to particular tissues (e.g., gut cells), and our whole-body measurements might not have been sensitive enough to pick up these subtle effects on the level of DNA methylation in specific tissues. In contrast, genes that appear to be both differentially expressed and differentially methylated were those with a rather general function, such as heat shock proteins and large ribosomal proteins. Genes encoding these proteins might perhaps show a more consistent expression across a range of cell types. Further analysis of DNA methylation in germ cells and differentiated cell types, and data on the relationship between DNA methylation and gene expression, could substantiate the breadth and stressor-specificity of DMPs in germ cells, their persistence during somatic cell differentiation, and functional relevance.

The epigenetic changes induced in offspring of exposed mothers commonly persisted until at least the fourth generation. The exception was the 5-azacytidin treatment, which demonstrates that modification of DNA methylation typically is restored from one generation to the next. Overall, only about 1% of sites that became hypomethylated in offspring of 5-azacytidin-exposed mothers remained hypomethylated in the F4 generation. This epigenetic resetting makes it the more striking that nearly half of the DMPs observed in the offspring of mothers exposed to microcystin, zinc, or high temperature actually persisted until the F4 generation. This suggests that naturally occurring stressors modify DNA methylation in a way that reliably allow those modifications to be passed on to subsequent generations in *Daphnia*. These results substantiate and extend previous work on pools of *Daphnia* individuals that indicated that methylation patterns induced upon salinity stress or gamma radiation can be detected until the F3 generation [32].

The mechanism by which DNA methylation is inherited remains poorly understood. Both direct copying of methylation states and involvement of small RNA molecules in RNA-directed DNA methylation [47] are potential mechanisms. MicroRNA expression in eggs of *Daphnia* vary as a result of maternal stress, but there is no evidence that differences in the expression of these RNAs persist for several generations [48]. However, this does not rule out that other forms of small RNAs, such as piRNA or tsRNA, are involved. RNA-mediated mechanisms may make it more likely that environmentally induced variation in DNA methylation will be inherited also during sexual reproduction and through both parents. More generally, establishing the mechanisms of transgenerational persistence of variation in DNA methylation will help to understand the extent to which it represents a flexible mechanism of inheritance that can contribute to ecological and evolutionary dynamics.

Despite a high incidence of parthenogenesis, *Daphnia* are famous for their ability to adapt rapidly to environmental stressors, including to all three naturally occurring stressors of this study (e.g., toxic cyanobacteria [49]; metal pollution [50], and high temperature [51]). Interestingly, such adaptations can be rapidly lost if conditions improve [50]. Laboratory studies have also demonstrated strong environmentally induced maternal effects (e.g., toxic cyanobacteria [52]; metal pollution [53]; temperature [54]), sometimes persisting for several generations [27, 55, 56]). The persistence of environmentally induced DNA methylation from one generation to the next that we demonstrate here could partly contribute to such transgenerational effects, and suggests that low genetic diversity may not prevent *Daphnia* populations from responding to selection. Thus, our results suggest that epigenetic inheritance can contribute to the adaptability of *Daphnia*, allowing populations to persist even under rapid and severe environmental change.

## Materials and Methods

### Daphnia husbandry and experimental design

A stock of *Daphnia magna* was sourced from Lake Bysjön (surface area 10 ha, 55°40′32″N 13°32′42″E) in Southern Sweden. Single clonal lines were kept under laboratory conditions [52] and allowed to reproduce asexually for 12 months before the onset of the experiment. All experiments in this study used a single clone to minimize any genetic effects. Applying the experimental design shown in Figure 1, individual *Daphnia* of the generation F0 were exposed to environmental stressors from birth to first reproduction (more detail in Table S1). Since the maternal treatment stopped before the egg cells were released into the brood pouch, the F1 generation was exposed as germ cells to the stressors but not as embryos [57]. Following the first brood, all individuals of the F0 generation were maintained under control conditions. We propagated these lines down to generation F4 by isolating five offspring from the second brood in each generation and keeping them under control conditions. Subsequent generations (F2, F3 and F4) did not encounter the stressors. We collected individuals of generations F1, F2 and F4 for whole-genome bisulfite sequencing directly after they produced their second brood (i.e., as adults). We omitted the F3 generation and used the F4 generation instead to gain insights into truly transgenerational effects on DNA methylation. This experiment was performed twice simultaneously (T1 and T2), to account for potential incompleteness due to the extinction of maternal lines, and the most complete experiment for each stressor was subjected to further analyses. The experiment T1 for 5-azacytidine, microcystin and high temperature, and experiment T2 was selected for the zinc treatment. Controls were matched within each of these two experimental groups (i.e., the effects of zinc were evaluated against its corresponding T2 control line, and the effects of the other three treatments against the T1 control line). Previous analyses of DNA methylation in *Daphnia* have relied on pools of individuals [32, e.g., 58], but to avoid confounding effects, we applied a newly developed low input methodology [59] that allowed us to sequence individual *Daphnia*.

### Reproductive output as a proxy of fitness effects

To assess how the stressor treatments affect the lifetime reproductive success of exposed individuals and their descendants, we collected the age of first and second reproduction (in days) and the sizes of the first and second brood for all individuals of the selected experiment (T1 or T2; including those individuals not selected for bisulfite sequencing). We estimated fitness by calculating, for each individual, the intrinsic rate of population increase *r* with a univariate root finding algorithm (*uniroot* in R) using the Euler equation [for details, see 52]. To test whether fitness varied by treatment and generation we estimated fitness for each treatment by generation in a nested multilevel model, with generation nested within treatment (in Stan 2.21.0 accessed from R with rstan 2.21.2). We present fitness effects as the difference between the fitness for a particular treatment by generation and the fitness of the control treatment of the same generation (with negative effects indicating a reduction of fitness compared to the control).

### WGBS, read mapping and extraction of methylation values

DNA was extracted from whole individual *Daphnia* samples using the DNeasy blood and tissue kit (Qiagen^TM^, Valencia, CA, USA) and DNA concentrations were estimated using a Qubit Fluorometer (ThermoFisher Scientific). Three individuals per experimental unit were initially processed, but units containing samples with low DNA concentrations were supplemented with a fourth back-up sample (i.e., for some units, four rather than three samples were processed). Extracted DNA samples were subjected to library preparation using the SPLAT protocol [59] with minor modifications. Adapter oligos were modified at all the 5’- and 3’-ends not involved in ligation to reduce adapter dimer formation. The following adapter oligos were used: 5’AmMC6/GACGTGTGCTCTTCCGATCTNNNNNN/3’AmMo, 5’Phos/AGATCGGAAGAGCACACGTC/3’AmMo, 5’AmMC6/ACACGACGCTCTTCCGATCT, and 5’AmMC6/NNNNNNAGATCGGAAGAGCGTCGTGT/3’AmMo. All oligos were purchased from IDT. Libraries were sequenced on six lanes of an Illumina HiSeqX instrument in randomized order. Sequencing data were processed within the framework of the nf-core methylseq workflow version 1.5 [60] (Figure S3). In summary, raw reads of 64 fastq files were trimmed of adapter sequences using Trim Galore! with default parameters. Trimmed reads were mapped to the *Daphnia magna* reference genome GCA_003990815.1 [genome size: 123 Mb; 61] using Bismark [62] with the paired-end setting and with parameter settings “-q --score-min L,0,-0.2 --ignore-quals --no-mixed --no-discordant --dovetail --maxins 500 --directional”. Cytosine methylation from deduplicated sequence data was generated using bismark_methylation_extractor [62] with parameter settings “--ignore_r2 2 --ignore_3prime_r2 2 --no_overlap”.

Six libraries were excluded from the analyses due to low read mapping rate and cytosine site coverage (<15% mapping rate, <1X mean coverage and <5X median coverage). Furthermore, four libraries were excluded on the basis of being PCA outliers in CpG percent methylation. Of these outliers, three were characterised by high percentage of methylated CpG (>97% percentile of the remaining libraries), which were at levels similar to the six libraries excluded due to low read mapping rates. In total, 54 libraries passed the quality control and proceeded to further analyses. These libraries had a mean coverage of 5.3X and a mean mapping rate of 48% (Table S2).

### Differential methylation analysis

The Bioconductor R package methylKit_1.12.0 [63] was used to carry out differential methylation analysis comparing each treatment (case) against untreated (control) groups. For each case and control selection, we only consider CpG sites with a minimum of 5 total read counts in all samples in all F-generations. The read counts of all sites that passed this filter were normalised by a library specific scaling factor as computed by a median coverage normalisation in methylKit. Furthermore, sites were filtered to consider only variable sites with sample standard deviations in percent methylation values of ≥0.5 (per case-vs-control group; see below). Overall, the initial set of >8 M CpG sites called per sample was reduced to an average of 2.8 M sites (range from 2.3-3.3 M) that were tested for differential methylation analysis.

Within the methylKit framework, we used the Wald test for hypothesis testing and beta binomial with overdispersion correction and parameter shrinkage to model the proportion of methylated CpG at a site. We quantitatively confirmed the main results using logistic regression models (Figure S4). The Benjamini-Hochberg method was used for multiple testing correction.

The case-vs-control statistical tests to identify differentially methylated CpG sites were carried out independently for each F-generation of case samples. However, to control for any generational epigenetic drift that could add to stochastic noise in the controls, the same set of all control samples across generations was used for each statistical test, i.e., 3 cases of F1 vs 9 controls (F1+F2+F4), 3 cases of F4 vs 9 controls (F1+F2+F4). Lastly, the statistical significance of differentially methylated CpG (DMP) sites was adjusted with a 5% FDR in each test.

### Identification of transgenerational DMPs

We defined transgenerational DMPs as being CpG sites that have acquired a treatment induced methylation state in the F1 generation (i.e., differentially methylated in the three F1 samples compared to the nine control samples), and for which methylation states are consistently maintained in the succeeding F2 and F4 generations (i.e., differentially methylated in both F2 vs control and F4 vs control). No minimum methylation difference was imposed. Furthermore, the statistical significance of DMP overlap across generations was obtained using the permutation function *permTest* in the R package regioneR_1.20.0 [64] and resampling randomly from all tested CpGs.

To robustly verify that the treatment-induced transgenerational DMPs are stably inherited across generations and not due to stochastic events, we used two additional strategies for data analyses. First, we carried out a permutation test by randomly assigning sample labels. For each selected treatment and control pair, we permuted their sample labels by shuffling case/control labels (e.g., zinc and control) and generation labels (i.e., F1, F2 and F4). We carried out 100 permutations of sample labels. After permuting sample labels, differential analysis was carried out as described above, i.e. 3-vs-9 per generation, with the same set of 9 “controls” in each generation. The null hypothesis was no association between CpG methylation and sample labels and we expect that randomly shuffling the sample labels would fulfil the null hypothesis. By inspecting quantile-quantile (Q-Q) plots (Figure S1 and S2), we compared the distribution of true labels with that of randomly shuffled labels and assessed if the former was associated with lower *P*-values and a higher number of transgenerational DMPs (i.e., significant DMPs shared across the three generations). Since this strategy does not mitigate the potential bias stemming from using the same set of control samples (i.e., all nine control samples) in each of the statistical tests, we also adopted a second strategy of identifying DMPs to exclude the possibility that this non-independence inflates the number of DMPs. To this end, we selected candidate environmentally induced CpGs by selecting outliers from comparisons of three cases versus three controls in the F1 generation using a lenient 20% FDR. We then tested these candidate CpGs and asked which of them also meet the criterion of being differentially methylated at a 5% un-adjusted *P*-value cut-off in the F2 and F4 generations by performing three cases versus three controls tests within these two generations. When comparing this alternative set of DMPs against the set obtained using the original 3-vs-9 approach, we found that between 77% and 25% of the original approach were also identified by the alternative approach (Table S15), with the alternative approach being generally more stringent (i.e., producing lower numbers of transgenerational DMPs).

### Annotation of DMPs and gene ontology analyses

We assigned each DMP to a nearest gene or a gene unit (i.e., exon or intron) by cross-referencing its genomic position with the GTF annotation from the reference assembly. This was carried out using BEDOPS closest-feature [65]. To obtain functional annotation such as gene ontology for the *Daphnia* genome, we used eggnog 5.0 [66] (emapper-2.1.2) with default parameters but restricting to the taxon Arthropoda. The enrichment analysis of GO terms was carried out using the R package topGO (version 2.40.0) and Fisher’s exact test.

### Cross-referencing differentially methylated genes with differentially expressed genes identified in the literature

To assess if hypo- or hyper-methylated genes in this study are those demonstrated to be differentially expressed upon exposure to a given stressor, we systematically screened the literature for relevant transcriptomic studies. We conducted a literature search using ISI *Web of Science* (v.5.30) with search terms specific to each dataset. We used the search terms ‘Daphnia’ and ‘transcriptomic*’, ‘RNAseq’, ‘gene expression’, ‘microarray’, along with one of the following: ‘*zinc*’, ‘microcystin*’, ‘temperature*’ or ‘*azacytidine*’. We excluded studies that used experimental designs that are too dissimilar from our settings (e.g., in terms of exposure duration). Quantitative comparisons between the set of differentially methylated genes identified in the present study and differentially expressed genes taken from the literature is hampered by a number of facts (e.g., differentially expressed genes are not reported in a standardized way, overrepresented GO terms are rarely reported, assigning gene orthologs between different *Daphnia* species, or assigning corresponding genes between different genome versions of the same species, is not straightforward). In addition, these studies often report up to 30% of all transcripts as differentially expressed, which precludes quantitative enrichment analyses. We therefore restricted our analysis to cross-referencing the key genes singled out in genome-wide, unbiased approaches against the differentially methylated genes identified in our study. We manually compared gene sets and regarded genes as shared when they are semantically highly similar. For example, we considered ‘heat shock protein 70 Bbb’ similar to ‘heat shock factor protein-like, transcript variant X7’.

## Acknowledgements

We thank Hanna Laakkonen for assistance with DNA extractions and Alexander Hegg for assistance with *Daphnia* experiments.

## Data availability

All sequences generated in this study have been deposited in NCBI Sequence Read Archive (SRA) with accession number PRJNA760269. Data for the fitness analyses are deposited in Dryad (*DOI to be generated during submission*).

## Code availability

Code for the analysis of DNA methylation is available on Bitbucket (https://bitbucket.org/scilifelab-lts/t_uller_1801/). Code for the fitness analyses is deposited in Zenodo (DOI: 10.5281/zenodo.5635792).

## Author contributions

R.R. and T.U. conceived the study; N.F. and T.U. coordinated the study; R.R., E.W.T, B.T.H. and T.U designed the study; R.R. performed the experiments; A.R. performed optimization of library preparation and generated the sequencing libraries; L.V. analysed the sequence data with input from N.F., R.R., M.R., B.N. and T.U.; R.R. analysed fitness data with input from N.F. and T.U.; N.F. and T.U. wrote the manuscript with input from L.V., R.R., E.W.T and B.T.H. All authors approved the final manuscript.

