## Supplementary Figures for "Environmentally-induced DNA methylation is inherited across generations in an aquatic keystone species (*Daphnia magna*)"

This file contains:

-4 supplementary figures

Supplementary Material

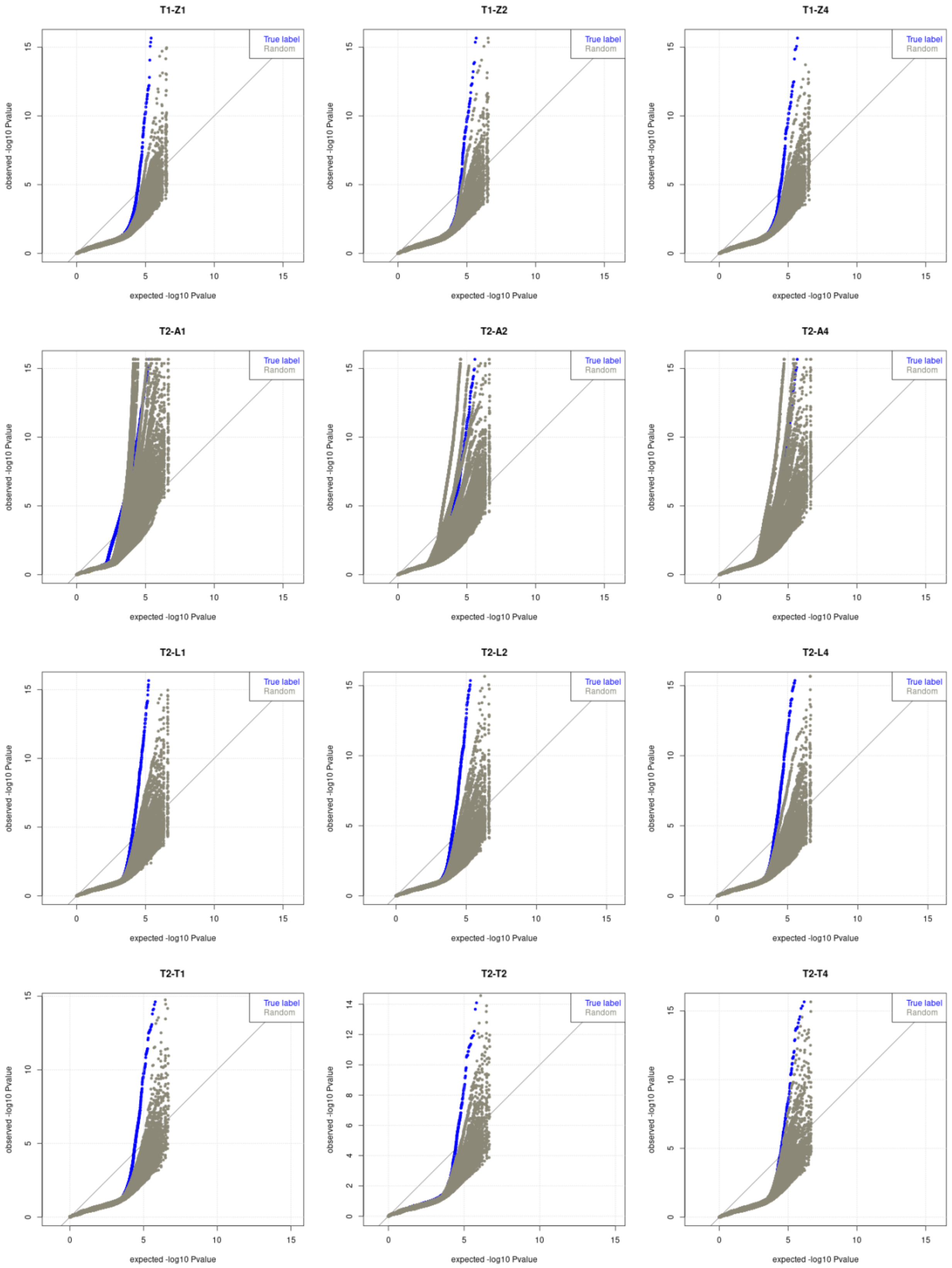

**Figure S1. Summary quantile-quantile (Q-Q) plot for the beta binomial model.** Q-Q plots show -log10 *P*-values from the null hypothesis on the x-axis (expected) and from a differential methylation analysis on the y-axis (observed). Note that under the null hypothesis, it is expected that *P*-values follow a uniform distribution. Q-Q plots were generated for each instance when true and random sample labels were used. If some of the tested CpGs deviate from the null hypothesis (i.e., statistically significant DMPs), we should see their *P*-values above the diagonal line. We expect that the statistical test using random labels will consistently result in a lower range of observed -log10 *P*-values than when the true labels are used.

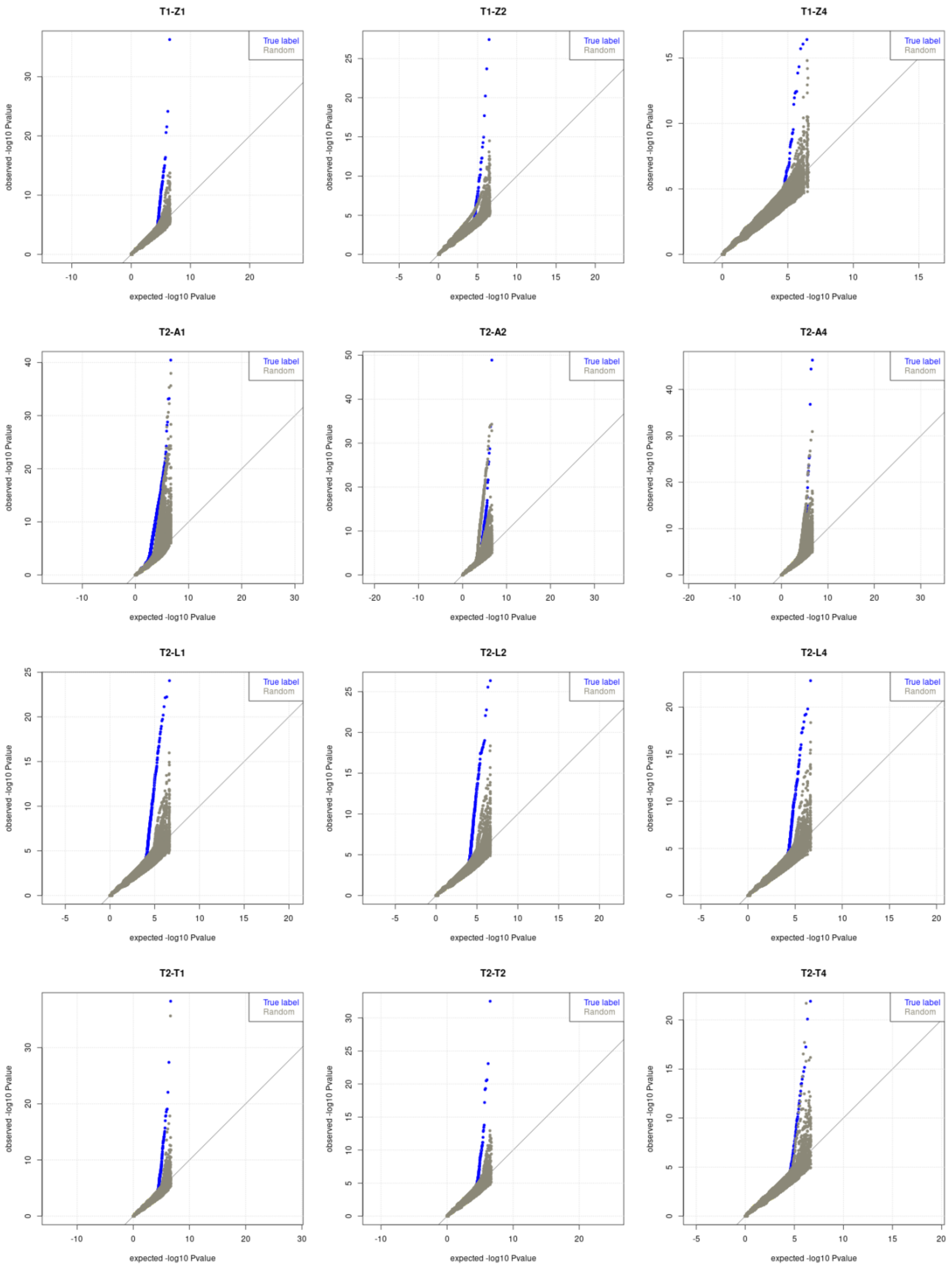
T1-Z1
T1-Z2
T1-Z4
T2-A1
T2-A2
T2-A4
T2-L1
T2-L2
T2-L4
T2-T1
T2-T2
T2-T4
True label
Random
observed -log10 Pvalue
expected -log10 Pvalue

**Figure S2. Summary quantile-quantile (Q-Q) plot for the logistic regression model.** Q-Q plots show -log10 *P*-values from the null hypothesis on the x-axis (expected) and from a differential methylation analysis on the y-axis (observed). Note that under the null hypothesis, it is expected that *P*-values follow a uniform distribution. Q-Q plots were generated for each instance when true and random sample labels were used. If some of the tested CpGs deviate from the null hypothesis (i.e., statistically significant DMPs), we should see their *P*-values above the diagonal line. We expect that the statistical test using random labels will consistently result in a lower range of observed -log10 *P*-values than when the true labels are used.

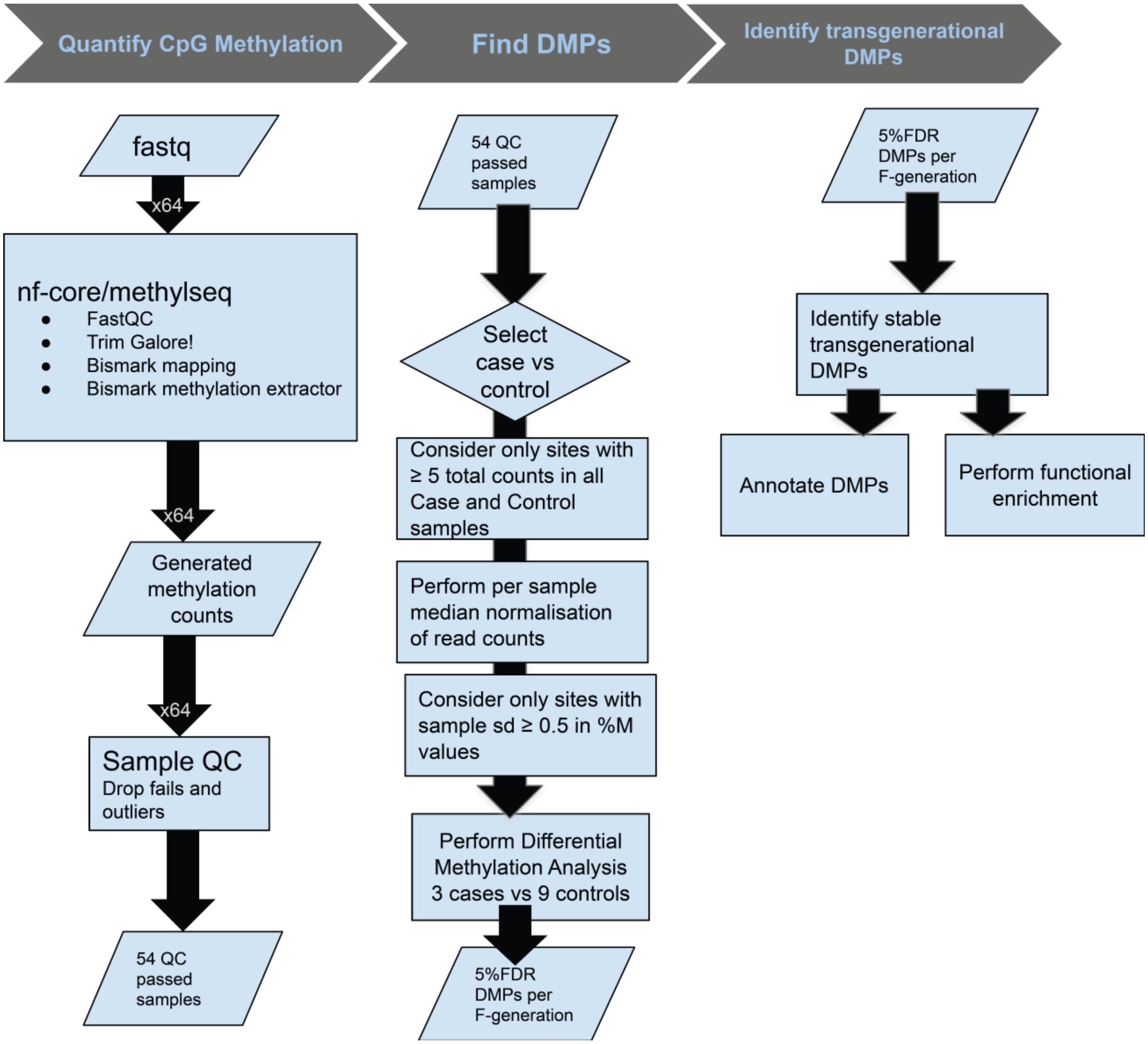


**Figure S3. Workflow of the bioinformatics analyses.** Schematic representation of the bioinformatics analysis that was employed to identify transgenerational DMPs from raw sequence reads derived from whole-genome bisulfite sequencing. Details are described in the Methods section.

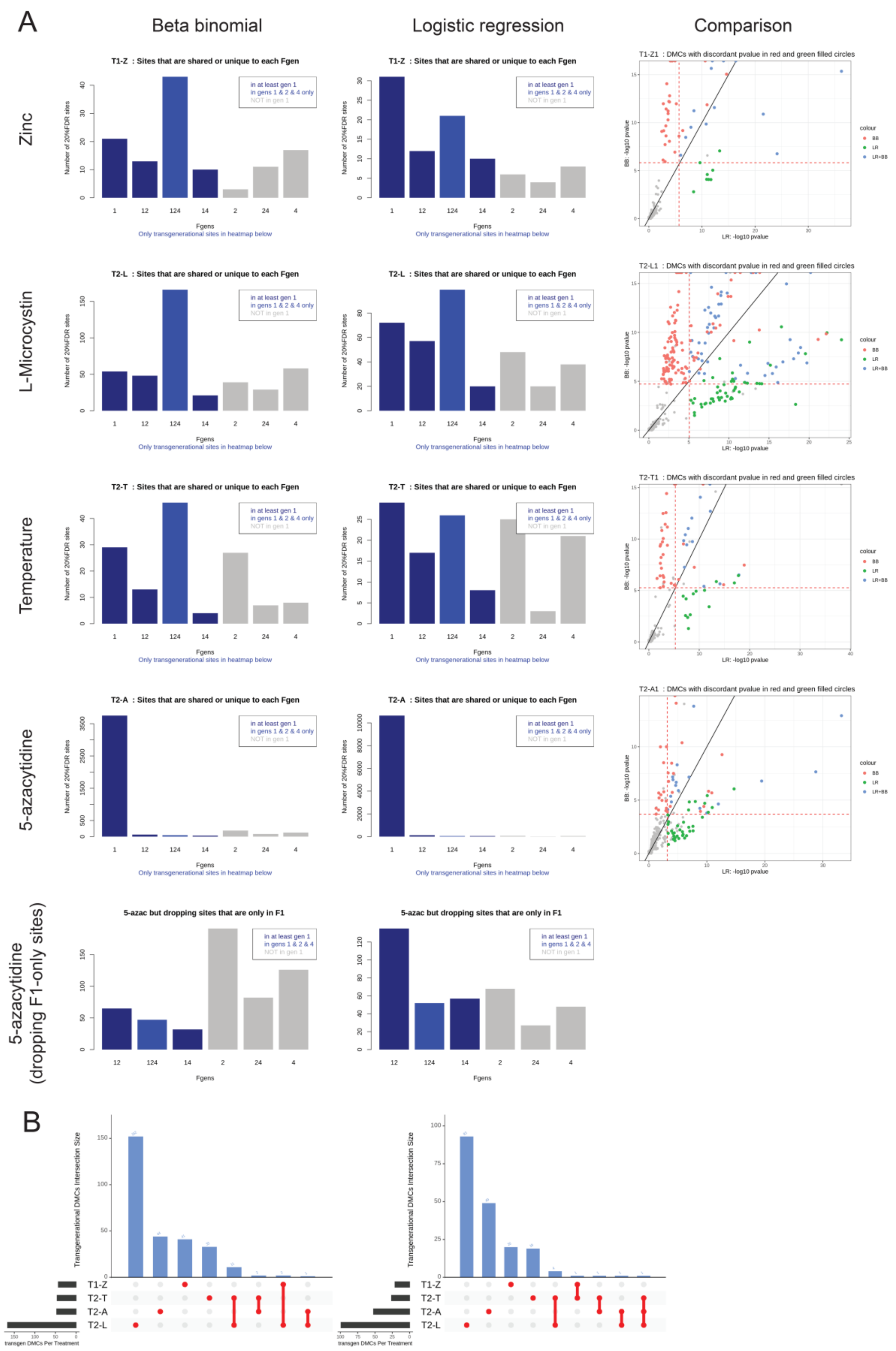
A
Beta binomial
Logistic regression
Comparison
Zinc
L-Microcystin
Temperature
5-azacytidine
5-azacytidine (dropping F1-only sites)
T1-Z : Sites that are shared or unique to each Fgen
T2-L : Sites that are shared or unique to each Fgen
T2-T : Sites that are shared or unique to each Fgen
T2-A : Sites that are shared or unique to each Fgen
5-azac but dropping sites that are only in F1
in at least gen 1
in gens 1 & 2 & 4 only
NOT in gen 1
in gens 1 & 2 & 4
Number of 20%FDR sites
Fgens
Only transgenerational sites in heatmap below
T1-Z1 : DMCs with discordant pvalue in red and green filled circles
T2-L1 : DMCs with discordant pvalue in red and green filled circles
T2-T1 : DMCs with discordant pvalue in red and green filled circles
T2-A1 : DMCs with discordant pvalue in red and green filled circles
BB: -log10 pvalue
LR: -log10 pvalue
colour
BB
LR
LR+BB
B
Transgenerational DMCs Intersection Size
T1-Z
T2-T
T2-A
T2-L
transgen DMCs Per Treatment

**Figure S4. Comparison of DMP overlap across generations between the beta binomial and logistic regression statistical modelling at a 20% FDR.** (A) Left column shows the number of DMPs per generation and overlap between generations for each stressor (rows) using a beta binomial model. Middle column show the same but using a logistic regression model. Right column shows -log10 *P*-values derived from the beta binomial modelling on the x-axis and the logistic regression modelling on the y-axis for the F1 generation of each stressor. Color-code highlights significant DMPs for each modelling approach. (B) Barplot presenting the overlap of transgenerational DMPs across the treatments for beta binomial (left) and logistic regression modelling (right). Horizontal bars represent the total number of transgenerational DMPs for each stressor, and vertical bars represent the DMPs unique to a given stressor (four bars to the left), or shared between two stressors (four bars to the right). No DMP was shared across all treatments, but between one and eleven are shared by two stressors. Note that these analyses have been conducted at a 20% FDR to increase the number of DMPs that could be compared between the two approaches.
